# A decoupled, modular and scriptable architecture for tools to curate data platforms

**DOI:** 10.1101/2020.09.28.282699

**Authors:** Moritz Langenstein, Henning Hermjakob, Manuel Bernal Llinares

## Abstract

**Motivation:** Curation is essential for any data platform to maintain the quality of the data it provides. Existing databases, which require maintenance, and the amount of newly published information that needs to be surveyed, are growing rapidly. More efficient curation is often vital to keep up with this growth, requiring modern curation tools. However, curation interfaces are often complex and difficult to further develop. Furthermore, opportunities for experimentation with curation workflows may be lost due to a lack of development resources, or a reluctance to change sensitive production systems.

**Results:** We propose a decoupled, modular and scriptable architecture to build curation tools on top of existing platforms. Instead of modifying the existing infrastructure, our architecture treats the existing platform as a black box and relies only on its public APIs and web application. As a decoupled program, the tool’s architecture gives more freedom to developers and curators. This added flexibility allows for quickly prototyping new curation workflows as well as adding all kinds of analysis around the data platform. The tool can also streamline and enhance the curator’s interaction with the web interface of the platform. We have implemented this design in cmd-iaso, a command-line curation tool for the identifiers.org registry.

**Availability:** The cmd-iaso curation tool is implemented in Python 3.7+ and supports Linux, macOS and Windows. Its source code and documentation are freely available from https://github.com/identifiers-org/cmd-iaso. It is also published as a Docker container at https://hub.docker.com/r/identifiersorg/cmd-iaso.

## 1 Introduction

The URLs to access data collections can change over time, leading to broken references. Many popular data collections are also accessible through more than one provider. The identifiers.org registry contains manually curated, high-quality metadata for hundreds of data collections, mainly from the Life Sciences domain (Juty *et al*., 2013). For each entry, it stores, amongst other metadata, a description of the data collection, a regular expression of the locally unique identifier pattern and the set of resources providing the data collection. For each provider, identifiers.org stores the data access endpoint of the resource. To facilitate the consistent and stable cross-referencing of data sets, identifiers.org also provides automatic resolution from globally unique compact identifiers to resource provider URLs (Wimalaratne *et al*., 2018).

Therefore, the primary curation objective of identifiers.org is to ensure that compact identifiers are resolved to the most reliable provider of the requested resource (Juty *et al*., 2013). While identifiers.org is a meta-registry of data platforms, the necessity for continuous curation extends to all providers of scientific information (Odell *et al*., 2017).

## 2 Methods of Curation

In order to provide reliable resolution of compact identifiers, identifiers.org must maintain accurate records and reliability scores for each resource provider. While there has long been an automatic regular availability check of provider endpoints (Juty *et al*., 2013), it has traditionally been based on HTTP status codes. A curator then still had to manually investigate the type of error, for example to distinguish between a planned maintenance outage and a failure of the provider.

We designed cmd-iaso to help with the current curation workflows, as well as any future ones. All curation workflows are split into information gathering, data analysis and augmented interactive curation. Currently, the tool implements the entire curation workflow for assessing the reliability of resource providers. Expensive data gathering and analysis can be pre-computed in the background. Additionally, cmd-iaso also supports analysis routines that can be performed live while the curator is interacting with the tool. The results of this analysis can then be presented to the curator through an augmented user interface of the existing platform.

## 3 Implementation

The curation problems, which cmd-iaso was designed to assist with, are challenging. Merely extending the identifiers.org platform would have limited experimentation with the design of the curation tool. During the development of cmd-iaso, extensive exploration of different approaches and iteration upon them was required.

cmd-iaso’s design is decoupled from identifiers.org, treats the platform as a black box and only communicates with it through its public APIs (see Fig. 1). It is written in the scripting language Python, which reduces the overhead for prototyping. This design decision simplifies implementing new curation pipelines. Technical knowledge of how the underlying data platform functions is not required. cmd-iaso is also designed as a modular tool, thereby supporting agile and iterative development. Please refer to supplementary material III and IV for details on the implementation of cmd-iaso’s modular plugin system as well as an example analysis using the tool, respectively. This system also allows individual plugins to utilise external data sources without affecting other workflows.

**Fig. 1.**
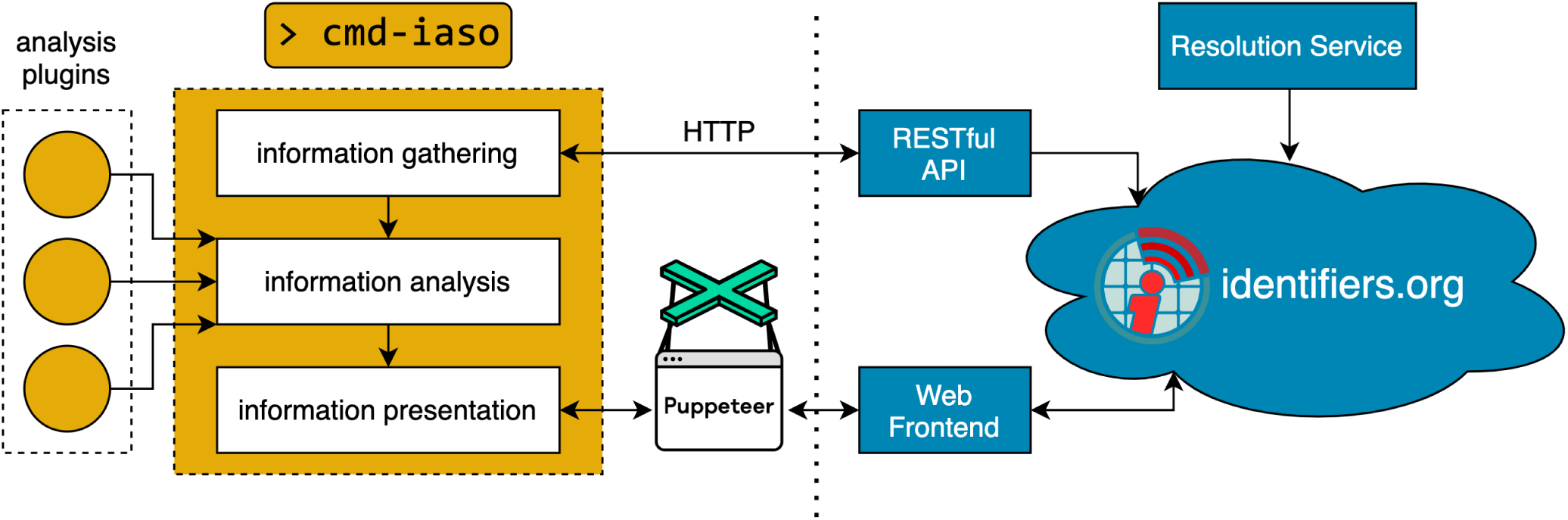
Software architecture of cmd-iaso.

The most significant feature of cmd-iaso is its interactive curation workflow, during which the curator is guided through the issues which have been identified. The tool can be run in a text-only terminal-based mode. However, cmd-iaso can best assist curation in its browser-based mode, in which it augments the existing web application of the platform. cmd-iaso uses pyppeteer, a Python port of the browser automation library Puppeteer. Puppeteer can launch or connect to a session of the Chrome browser and take full control over it (see Fig. 1). For instance, the library can inject new elements and information into any website and interact with its existing contents.

If cmd-iaso is run in its browser-based curation mode, it injects a control interface into identifiers.org’s website to allow the curator to jump between the entries identified for curation quickly. It also automatically navigates to the corresponding page in the registry and augments it with an information overlay. This overlay contains information about the issue, hyperlinks to referenced websites as well as any proposed corrections. For a visualisation of a typical curation session using cmd-iaso as well as more implementation details, please see supplementary materials I and II, respectively. It is worth emphasising that the entire augmentation only occurs locally in the curator’s browser. This augmentation is the perfect example of how a decoupled tool can extend and improve upon the existing interaction between the curator and their platform.

## 4 Discussion

We have proposed a decoupled, modular and scriptable architecture for a curation tool, which opens up the possibility for agile development and a diverse, easily maintainable set of plugins. cmd-iaso currently only uses a flexible plugin system in the data analysis stage. If this design were applied to the entire tool, cmd-iaso could become a general and highly customisable curation toolbox. It could then support the integration and curation of different data platforms with various analysis methods.

This approach of designing a decoupled, modular and scriptable curation tool can already be applied to any data platform. Its design principles could even be transformed into operational guidelines, as they make this architecture very suitable to close collaboration between curators and developers. The design also envisions a modern interpretation of the role of curators, in which they have an increasing ownership of and responsibility for the tools that support their curation workflows. cmd-iaso’s design allows for curators to be better equipped for the rapidly changing needs and magnitude of data in Life Sciences today (Tang *et al*., 2019).

## Supporting information

Supplementary Material III

## Supplementary information

Supplementary materials are available at *bioRxiv* online.

