## Supplementary Material I for "A decoupled, modular and scriptable architecture for tools to curate data platforms"

A decoupled, modular and scriptable architecture for tools to curate data platforms
Supplementary Material I

Augmenting an Existing Web Application with Puppeteer: Demonstration of the curation tool cmd-iaso

https://youtu.be/9CwODVae000

Moritz Langenstein, Henning Hermjakob and Manuel Bernal Llinares
