## Supplementary Material II for "A decoupled, modular and scriptable architecture for tools to curate data platforms"

### Augmenting an Existing Web Application using Puppeteer

Moritz Langenstein, Henning Hermjakob and Manuel Bernal Llinares

September 28, 2020

The most significant feature of cmd-iaso is its interactive curation workflow, during which the curator is guided through the issues which have been identified. cmd-iaso uses pyppeteer, a Python port of the browser automation library Puppeteer. It was originally designed to automate integration tests of web application user interfaces. Puppeteer can launch or connect to a session of the Chrome browser and take full control over it. For instance, the library can inject new elements and information into any website and interact with its existing contents.

You can launch a new or connect to a running Chrome browser instance using:

```python
# Launch a new instance of the Chrome browser
browser = await pyppeteer.launch(headless=False, ...)
# OR
# Connect to an existing instance of the Chrome browser
browser = await pyppeteer.connect(
    browserWSEndpoint=wsEndpoint, ...
)

page = await browser.newPage()
```

We have split the interaction of the curator with the augmented user interface into three parts. First is the navigator, which simply navigates the Chrome browser to the most relevant page of the existing web platform using `await page.goto(url)` (please refer to the pyppeteer API) in the most simple form.

Secondly, there is the controller that provides a control interface to the curator to jump between issues. To provide the additional interface, we inject CSS and a JavaScript script that adds the controller HTML if it has not been inserted yet.

For instance, with a custom `PyppeteerCoordinator` (please refer to cmd-iaso's source code) that takes care of injection (based on `page.evaluate(script: str, *args)` – please refer to the pyppeteer API), we create the controller interface using:

```python
async with PyppeteerCoordinator(page) as coordinator:
    with suppress(pyppeteer.errors.NetworkError):
        await coordinator.addStyleTagWithId(
            "header.css", "iaso-header-style"
        )

        await coordinator.evaluateScript("header.js")

        await coordinator.evaluateScript(
            "controller.js", buttons
        )
```

Here header.js creates a curation-only layer with the HTML Element ID `iaso-header` in which controller.js inserts a sidebar with control buttons:

```javascript
function (buttons) {
    let header = document.getElementById("iaso-header");

    // Check that the header has already been created
    if (header === null) {
        return;
    }

    header = header.firstElementChild;

    for (const [id, title] of buttons) {
        let button = document.getElementById(
            `iaso-controller-${id}`
        );

        // Only create the button if it is still missing
        if (button === null) {
            button = document.createElement("button");
            button.id = `iaso-controller-${id}`;
            button.onclick = () => {
                console.info(`iaso-controller-${id}`);
            };
            button.innerText = title;
        }

        // Appending all buttons will restore their order
        // even if they are created out of order
        header.appendChild(button);
    }
}
```

We use `console.info(msg)` to communicate any button press back to the Python script, which listens for console events using:

```python
def onconsole(console):
    if (console.type == "info" and
        console.text.startswith("iaso-controller-")
       ):
        print(f"The control button '{console.text[16:]}'" +
              " was pressed")

page.on("console", onconsole)
```

Last but not least, there is the informant/presenter which injects a separate information overlay into the interface using the same techniques showcased above.

The following screenshots will highlight the control interface (see Fig. 1) and information overlay (see Fig. 2) injected by cmd-iaso.

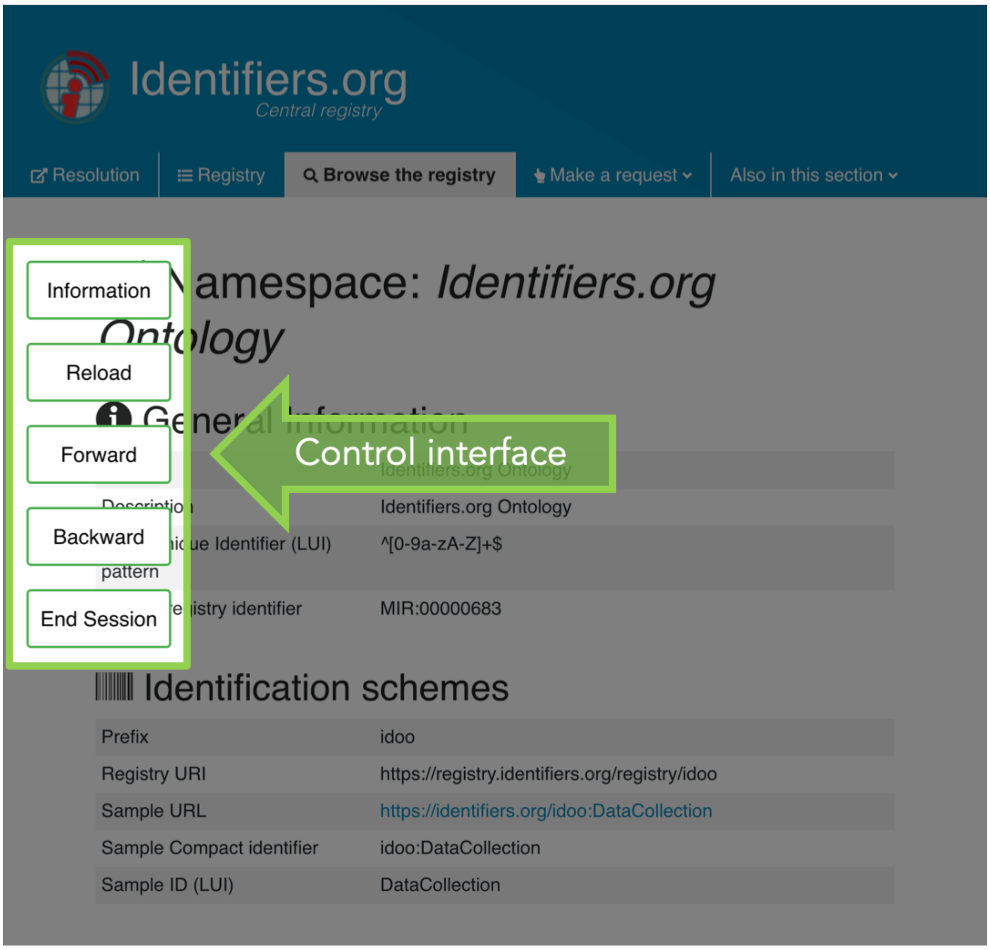


Figure 1: Example screenshot of the control interface provided by cmd-iaso on top of identifiers.org

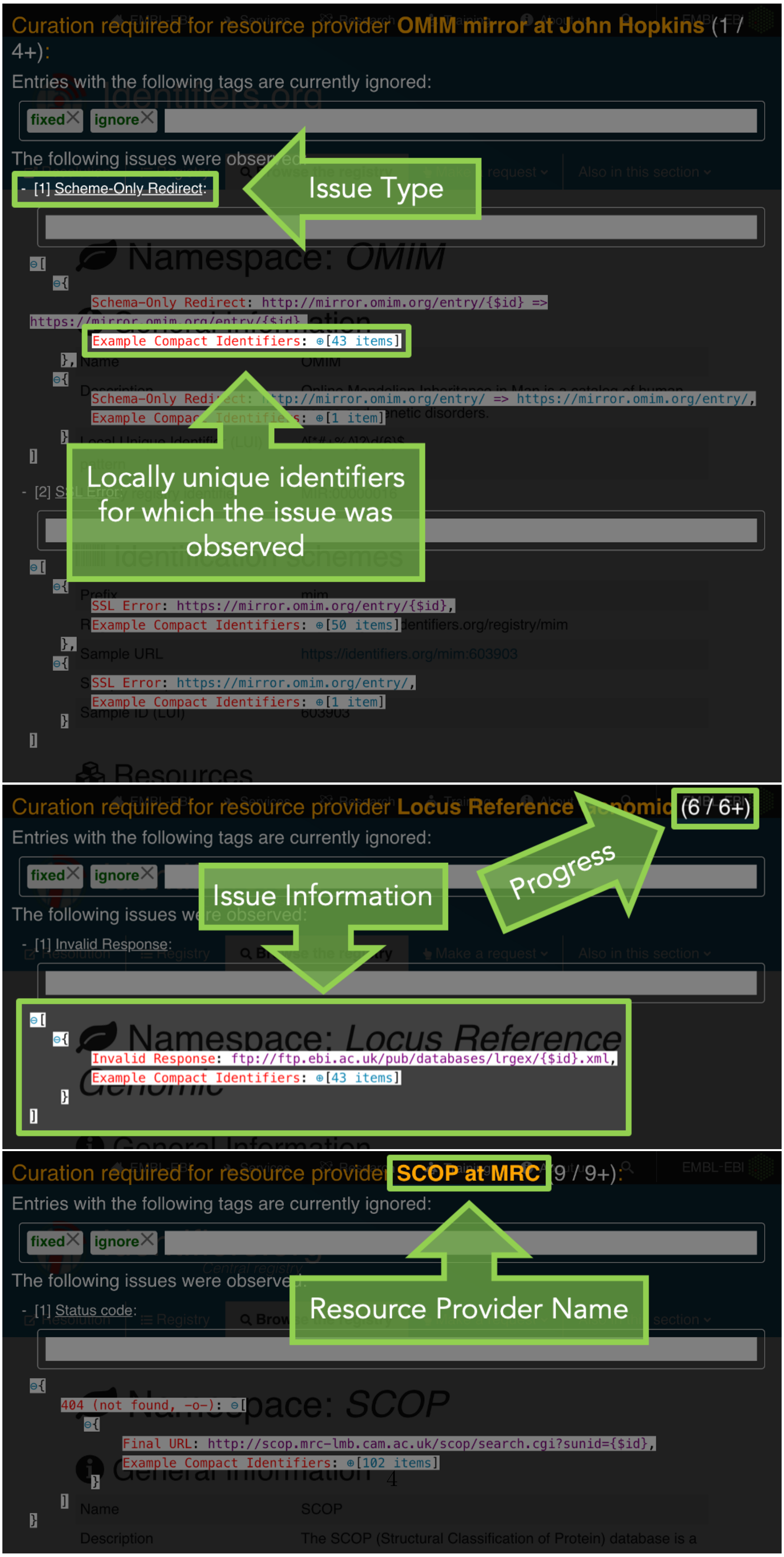
Curation required for resource provider OMIM mirror at John Hopkins (1 / 4+):
Entries with the following tags are currently ignored:
fixed ignore
The following issues were observed:
- [1] Scheme-Only Redirect:
Issue Type
Schema-Only Redirect: http://mirror.omim.org/entry/{$id} => https://mirror.omim.org/entry/{$id}
Example Compact Identifiers: [43 items]
Schema-Only Redirect: http://mirror.omim.org/entry/ => https://mirror.omim.org/entry/,
Example Compact Identifiers: [1 item]
Locally unique identifiers for which the issue was observed
- [2] SSL Error:
SSL Error: https://mirror.omim.org/entry/{$id},
Example Compact Identifiers: [50 items]
SSL Error: https://mirror.omim.org/entry/,
Example Compact Identifiers: [1 item]
Curation required for resource provider Locus Reference Genomic (6 / 6+)
Entries with the following tags are currently ignored:
fixed ignore
The following issues were observed:
- [1] Invalid Response:
Issue Information
Progress
Invalid Response: ftp://ftp.ebi.ac.uk/pub/databases/lrgex/{$id}.xml,
Example Compact Identifiers: [43 items]
Curation required for resource provider SCOP at MRC (9 / 9+):
Entries with the following tags are currently ignored:
fixed ignore
The following issues were observed:
- [1] Status code:
Resource Provider Name
404 (not found, -o-):
Final URL: http://scop.mrc-lmb.cam.ac.uk/scop/search.cgi?sunid={$id},
Example Compact Identifiers: [102 items]

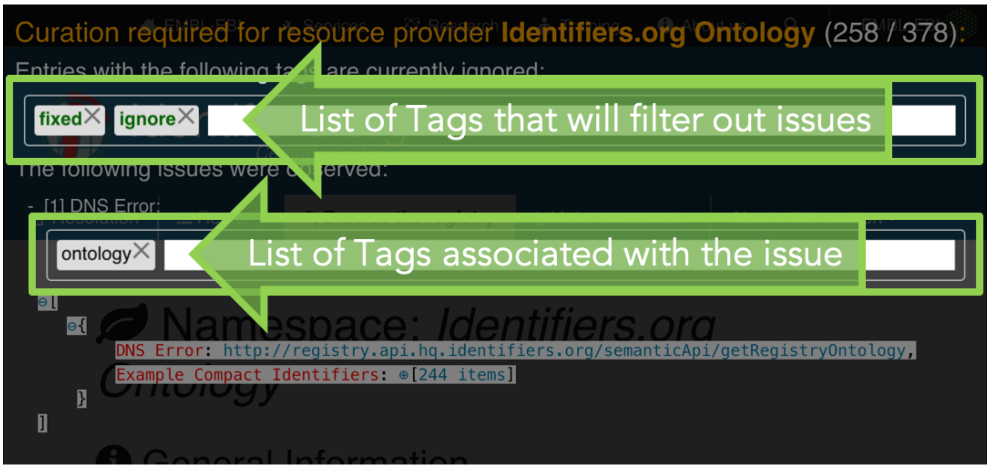


Figure 2: Example screenshots of the curation overlay provided by cmd-iaso on top of identifiers.org
