## Supplementary Material III for "A decoupled, modular and scriptable architecture for tools to curate data platforms"

A decoupled, modular and scriptable architecture for tools to curate data platforms
Supplementary Material III

### Modular Python Plugins for cmd-iaso

Moritz Langenstein, Henning Hermjakob and Manuel Bernal Llinares

September 28, 2020

cmd-iaso is designed as a modular command-line tool. Different data analysis strategies are provided by small plugin modules which can also be used to validate or filter out information that is irrelevant to the curator. While cmd-iaso provides some builtin default validators, the curator can very easily add new functionality. An abstract Python class is used to define the interface which the modules are required to implement. In cmd-iaso, we have created the following `CurationValidator` interface for this purpose:

```
from abc import ABC, abstractmethod
from typing import Union

class CurationValidator(ABC):
    @staticmethod
    @abstractmethod
    def check_and_create(data_entry) -> Union[
        CurationValidator, bool
    ]:
        """
        Returns False iff this data_entry cannot be included
         during curation at all.
        Returns True iff this validator has found nothing to
         report on this data_entry.
        Returns an instance of the CurationValidator subclass
         iff it found an issue to report for this data_entry.
        """
        pass

    @abstractmethod
    def present(self, presenter: CurationPresenter) -> None:
        pass
```

The plugins are registered as Python entry points under the public module path `curation_tool.plugins`. In our design, we require that a plugin is a non-abstract subclass, e.g. `MyValidator`, which implements the `CurationValidator` interface. Previously, entry points had to be registered with a packaging system like Flit, Poetry or Setuptools. Since PEP 621, entry points can be also be registered in the standardised `pyproject.toml` file:

```
[project]
name = "My Curation Plugin"

[project.entry-points."curation_tool.plugins"]
my_plugin = "my_module:my_validator:MyValidator"
```

The plugins can then be dynamically loaded into the tool as requested by the curator's selection using command line options. More generally, here is how you can iterate over all plugins registered under the `curation_tool.plugins` path:

```
try:
    from importlib.metadata import entry_points, EntryPoint

    def iter_entry_points(path):
        return entry_points().get(path, [])
except ImportError:
    # Running on pre-3.8 Python
    from pkg_resources import iter_entry_points, EntryPoint

for entry_point in iter_entry_points("curation_tool.plugins"):
    Validator = entry_point.load()

    if not issubclass(Validator, CurationValidator):
        # Validator is not a subclass of CurationValidator
        continue

    if len(Validator.__abstractmethods__) > 0:
        # Validator is not a non-abstract subclass of
        #  CurationValidator
        continue

    print(f"Successfully loaded {entry_point.name} into " +
          f"{Validator}")
```

This system is an example of how curators and developers can easily extend the command-line tool without having to modify its core source code.
