## Supplementary Material IV for "A decoupled, modular and scriptable architecture for tools to curate data platforms"

### Analysing the Reliability of Bioinformatics Resource Providers listed in identifiers.org using cmd-iaso

Moritz Langenstein, Henning Hermjakob and Manuel Bernal Llinares

September 29, 2020

cmd-iaso is a command-line tool to help curate the identifiers.org registry. In this Jupyter Notebook, we will go over the analysis one curation workflow provides in detail. You can also run this Jupyter Notebook online using:

launch binder

The issues, which we will identify and discuss below, were observed on 18/09/2020. Please note that we plan to share them with the external site operator, and update the identifiers.org records to ensure reliable URL resolution. Therefore, these issues will have likely have been resolved by the time of the publication of this work.

#### 1 Installation and Setup

First, we need to set up `cmd-iaso`. We will clone its source code from GitHub and install it in a fresh Python virtual environment `venv`.

[1]:
```
!git clone https://github.com/identifiers-org/cmd-iaso.git
!pip install virtualenv
!virtualenv venv
!venv/bin/pip install --upgrade pip
!venv/bin/pip install cmd-iaso/
```

For this analysis, we will import some standard library Python modules and define a JSON pretty-printing function which plugins into functionality used inside `cmd-iaso` itself.

[2]:
```
import gzip
import json
import pickle
import shlex
import urllib.request

def print_json(obj):
    code = shlex.quote(
        "from iaso.format_json import format_json; "
        + f"print(format_json({repr(obj)}, process_links=False))"
    )

    !echo {code} | venv/bin/python3
```

#### 2 Information collection

In this analysis, we will demonstrate the workflow to assess the reliability of resource providers to which the identifier.org platform redirects. For clarity, we will focus on one resource provider as an example: JWS Online Model Repository at Amsterdam.

Internally, this provider has been assigned the id `416`. We can use identifiers.org's API to examine the information the registry contains.

[3]:
```
with urllib.request.urlopen(
    'https://registry.api.identifiers.org/restApi/resources/416'
) as response:
    print_json(json.loads(response.read()))
```


```
{
  mirId: MIR:00100169,
  urlPattern: http://jjj.bio.vu.nl/models/{$id}/,
  name: JWS Online Model Repository at Amsterdam,
  description: JWS Online Model Repository at Amsterdam,
  official: False,
  providerCode: CURATOR_REVIEW,
  sampleId: curien,
  resourceHomeUrl: http://jjj.bio.vu.nl/models/,
  created: 2019-06-11T14:16:08.181+0000,
  modified: 2020-08-24T18:08:55.470+0000,
  deprecated: False,
  deprecationDate: None,
  _links: {
    self: {
      href: https://registry.api.identifiers.org/restApi/resources/416
    },
    resource: {
      href: https://registry.api.identifiers.org/restApi/resources/416
    },
    institution: {
      href: https://registry.api.identifiers.org/restApi/resources/416/
            institution
    },
    namespace: {
      href: https://registry.api.identifiers.org/restApi/resources/416/namespace
    },
    location: {
      href: https://registry.api.identifiers.org/restApi/resources/416/location
    },
    contactPerson: {
      href: https://registry.api.identifiers.org/restApi/resources/416/
            contactPerson
    }
  }
}
```

To assess the reliability of a resource provider, we need to test out how it responds to HTTP requests automatically. Therefore, we are most interested in the provider's `urlPattern` and `sampleId`. By replacing `{id}` in the `urlPattern` with any LUI (locally unique identifier) that is invalid in the resource's namespace, such as the `sampleId`, we get a URL which we can ping to check if the provider responds as expected.

In production, we would use `> cmd-iaso jobs jobs.json` to generate a list of such URLs to ping automatically. `cmd-iaso` would then take care to combine user-provided and randomly generated LUIs to cover a breadth of the space of identifiers of the namespace. It is important to note that `cmd-iaso` only generates random LUIs according to the resource's namespace's LUI regular expression `pattern`.

For this example, we will manually create this list consisting of the registered example ID `curien` and the randomly generated ID `7d_`. Both of these conform to the resource's namespace's LUI regex `pattern`, here `^\w+$`. Each job in the list consists of its resource provider ID, here `416`, the LUI, whether the LUI is random, and the full data access URL. We will write this list to the `jobs.json` file.

[4]:
```python
with open('jobs.json', 'w') as file:
    json.dump([
        (416, 'curien', False, 'http://jjj.bio.vu.nl/models/curien'),
        (416, '7d_', True, 'http://jjj.bio.vu.nl/models/7d_')
    ], file)
```

Next, we will create a folder `dump` for the `cmd-iaso` to store the scraping dumps in. We can now invoke the `> cmd-iaso scrape jobs.json dump` command.

[5]:
```
!mkdir dump -p
!echo 'y' | venv/bin/cmd-iaso scrape jobs.json dump
```

```
Loading the scraping jobs from jobs.json ...
Serving HTTPS Proxy on 0.0.0.0:34801 ...
100%|██████████| 2/2 [00:06<00:00, 3.08s/it, workers=0, processes=10]
```

#### 3 Data analysis

During scraping, `cmd-iaso` collects metadata about each response as well as, if possible, the content of the response. At the time of writing, this particular resource provider used TLS 1.0 encryption for their SSL connection, which is considered to be insecure today. If we run the scraping on an operating system which still allows HTTPS connections using TLS 1.0, `cmd-iaso` will be able to both flag the SSL error and record the response. Otherwise, it will critically fail during the ping and only report the SSL error. We can now take a look at the two raw dumps to see which is the case.

[6]:
```python
with gzip.open('dump/pings_416.gz', 'rb') as file:
    print_json(pickle.load(file))
    print_json(pickle.load(file))
```


```
{
  lui: curien,
```

```
  random: False,
  date: 2020-09-18 16:42:24,
  redirects: [
    {
      url: http://jjj.bio.vu.nl/models/curien,
      ip_port: 130.37.96.76:80,
      response_time: 66,
      status: 301,
      dns_error: False,
      ssl_error: False,
      invalid_response: False
    },
    {
      url: https://jjj.bio.vu.nl/models/curien,
      ip_port: None,
      response_time: None,
      status: None,
      dns_error: False,
      ssl_error: True,
      invalid_response: False
    }
  ],
  content: None,
  content-type: None
}
{
  lui: 7d_,
  random: True,
  date: 2020-09-18 16:42:24,
  redirects: [
    {
      url: http://jjj.bio.vu.nl/models/7d_,
      ip_port: 130.37.96.76:80,
      response_time: 62,
      status: 301,
      dns_error: False,
      ssl_error: False,
      invalid_response: False
    },
    {
      url: https://jjj.bio.vu.nl/models/7d_,
      ip_port: None,
      response_time: None,
      status: None,
      dns_error: False,
      ssl_error: True,
      invalid_response: False
    }
  ],
  content: None,
  content-type: None
```

```
}
```

`cmd-iaso` will now analyse and compact the raw contents of these data dumps into one structured findings file. For simple analysis based solely on the response metadata, `cmd-iaso` merely forwards the raw metadata. However, `cmd-iaso` also provides a more complicated analysis of the textual content of each response. You can read more about this `athena` analysis in `cmd-iaso`'s README.

[7]:
```
!echo 'y' | venv/bin/cmd-iaso dump2datamine dump datamine.json
```

```
Combining scraping dumps:   0%|          | 0/3 [00:00<?, ?it/s]
Loading scraped resource: 0it [00:00, ?it/s]
Loading scraped resource: : 0it [00:00, ?it/s]
Loading scraped resource:   0%|          | 0/1 [00:00<?, ?it/s]
Loading scraped resource:   0%|          | 0/1 [00:00<?, ?it/s]
Loading scraped resource:   0%|          | 0/1 [00:00<?, ?it/s]
Loading scraped resource:   0%|          | 0/1 [00:00<?, ?it/s]
Loading scraped resource:   0%|          | 0/1 [00:00<?, ?it/s]
The scraping DUMP at dump was successfully converted into a DATAMINE file at
datamine.json.
Loading scraped resource:   0%|          | 0/1 [00:00<?, ?it/s]
Combining scraping dumps:  33%|█         | 1/3 [00:00<00:00,
                                             172.88it/s]
```

This datamine file looks almost the same as the combination of the raw dumps but is missing the response contents. It also contains information about the environment in which we ran the scraping.

[8]:
```
with open('datamine.json', 'r') as file:
    print_json(json.load(file))
```


```
{
  environment: {
    machine: idorgdev,
    os: Linux-4.19.0-8-cloud-amd64-x86_64-with-debian-10.3,
    cpu: GenuineIntel Intel Core Processor (Skylake, IBRS) 6.94.3,
    cores: 1 x 8,
    memory: 4.53GiB,
    storage: 1.52GiB,
    cmd: scrape jobs.json dump --proxy launch --workers 32 --timeout 30
  },
  providers: [
    {
      id: 416,
      pings: [
        {
          lui: curien,
          random: False,
          date: 2020-09-18 16:42:24,
          redirects: [
            {
              url: http://jjj.bio.vu.nl/models/curien,
```

```
          ip_port: 130.37.96.76:80,
          response_time: 66,
          status: 301,
          dns_error: False,
          ssl_error: False,
          invalid_response: False
        },
        {
          url: https://jjj.bio.vu.nl/models/curien,
          ip_port: None,
          response_time: None,
          status: None,
          dns_error: False,
          ssl_error: True,
          invalid_response: False
        }
      ],
      empty_content: True
    },
    {
      lui: 7d_,
      random: True,
      date: 2020-09-18 16:42:24,
      redirects: [
        {
          url: http://jjj.bio.vu.nl/models/7d_,
          ip_port: 130.37.96.76:80,
          response_time: 62,
          status: 301,
          dns_error: False,
          ssl_error: False,
          invalid_response: False
        },
        {
          url: https://jjj.bio.vu.nl/models/7d_,
          ip_port: None,
          response_time: None,
          status: None,
          dns_error: False,
          ssl_error: True,
          invalid_response: False
        }
      ],
      empty_content: True
    }
  ],
  analysis: None
 }
]
}
```

#### 4 'Interactive' Curation

We can now use `cmd-iaso`'s interactive curation to walk us through the issues it has identified. We will use the `scheme-only-redirect`, `ssl-error` and `http-status-error` validators. The first validator finds HTTP redirects which only change the schema but not the rest of the URL. These extraneous redirects could potentially be avoided by updating the `urlPattern` stored in identifiers.org's registry. The second and third validators look for SSL and HTTP error codes in the responses, respectively. You can find their implementations on GitHub.

Firstly, we will make use of the `statistics` curation mode, which counts each error type and can give us a quick overall summary:

[9]:
```
!echo 'y' | venv/bin/cmd-iaso curate --statistics start resources
  datamine.json --validate scheme-only-redirect --validate ssl-error
  --validate http-status-error --random-luis-threshold=0 --discard-session
```

```
Loading the datamine file from datamine.json ...
tags.gz does not exist yet. Do you want to start with a new cross-session tags
store? [y/N]: Loading the identifiers.org registry ...
The data loaded was collected in the following environment:
{
  machine: idorgdev,
  os: Linux-4.19.0-8-cloud-amd64-x86_64-with-debian-10.3,
  cpu: GenuineIntel Intel Core Processor (Skylake, IBRS) 6.94.3,
  cores: 1 x 8,
  memory: 4.53GiB,
  storage: 1.52GiB,
  cmd: scrape jobs.json dump --proxy launch --workers 32 --timeout 30
}
The http-status-error, scheme-only-redirect and ssl-error validators were
loaded.
Starting the curation process of 1 entries ...
============================= Curation Statistics ===============================

In response to the current settings, 2 entries were identified for curation,
none of which were ignored because of their issues' tags.

The following issue types were identified for curation:
- SchemaRedirectError: 2 (0 ignored)
- SSLError: 2 (0 ignored)

================================================================================
Finishing the curation process ...
The current curation session has been discarded.
```

To find out about the details of each issue, we need to use the interactive curation mode `cmd-iaso` provides. Due to the small scope of this example, we only have one resource provider to curate. Therefore, we will tell `cmd-iaso` to end the curation session after displaying the first (and only) provider in need of curation. For a full visual demonstration of the curation in action, please refer to Supplementary Material I. Please note that we are using the text-only terminal curation mode here.

[10]:
```
!echo 'end' | venv/bin/cmd-iaso curate --controller terminal --navigator
  terminal --informant terminal start resources datamine.json --validate
  scheme-only-redirect --validate ssl-error --validate http-status-error
  --random-luis-threshold=0 --discard-session
```

```
Loading the datamine file from datamine.json ...
Loading the identifiers.org registry ...
The data loaded was collected in the following environment:
{
  machine: idorgdev,
  os: Linux-4.19.0-8-cloud-amd64-x86_64-with-debian-10.3,
  cpu: GenuineIntel Intel Core Processor (Skylake, IBRS) 6.94.3,
  cores: 1 x 8,
  memory: 4.53GiB,
  storage: 1.52GiB,
  cmd: scrape jobs.json dump --proxy launch --workers 32 --timeout 30
}
The http-status-error, ssl-error and scheme-only-redirect validators were
loaded.
Starting the curation process of 1 entries ...
             >>> https://registry.identifiers.org/registry/jws <<<
==================================== 1 / 1 ====================================
Curation required for resource provider JWS Online Model Repository at
Amsterdam:
The following issues were observed:
- [1] SSL Error: [
    {
      SSL Error: https://jjj.bio.vu.nl/models/{$id},
      Example Compact Identifiers: [
        1/1 x jws:curien (non-random),
        1/1 x jws:7d_ (random)
      ]
    }
  ] tagged with []
- [2] Scheme-Only Redirect: [
    {
      Schema-Only Redirect: http://jjj.bio.vu.nl/models/{$id} =>
                            https://jjj.bio.vu.nl/models/{$id},
      Example Compact Identifiers: [
        1/1 x jws:curien (non-random),
        1/1 x jws:7d_ (random)
      ]
    }
  ] tagged with []
==================================== 1 / 1 ====================================
Continue curation (rl, fw, bw, end, tags, ignore) [fw]: Finishing the
curation process ...
The current curation session has been discarded.
```

The issues, which we have identified and discussed above, were observed on 18/09/2020. Please note that we plan to share them with the external site operator, and update the identifiers.org records to ensure reliable URL resolution. Therefore, these issues will have likely have been resolved by the time of the publication of this work.

In this short example analysis, we have presented a detailed look at how `cmd-iaso` collects and analyses responses from resource providers to allow the curator to assess their reliability. If you would like to know more about the validator plugin system, which allows the curator to extend the analysis capabilities of identifiers.org easily, please refer to Supplementary Material III.
